# Pseudoreplication and inappropriate statistical tests in the analysis of preference index data from the T-maze group assay for *Drosophila behaviour*

**DOI:** 10.1101/2023.12.15.571933

**Authors:** Michael Winklhofer

**Affiliations:** Carl von Ossietzky Universität Oldenburg, School of Mathematics and Science, Institute for Biology and Environmental Science and Research Center Neurosensory Science, Oldenburg, Germany

## Abstract

The binary choice chamber (T-maze) assay has been used as a standard behavioural screen in Drosophila research to test mutants in terms of sensory discrimination skills and synaptic plasticity during memory consolidation and decay. Typically, ca. 100 individuals are tested as a group at the same time and the behavioural readout consists in counting the number of individuals in the testing tube exposed to a stimulus versus the number of flies in the control tube, with the normalized difference in fly count being defined as the batch preference index (PI). Unfortunately, the batch PI has widely been taken as a precise metric of group behaviour, to the point where ANOVA/t-tests have been considered the most powerful statistical tests for analyzing samples in terms of batch PI values, which has led to a hyperinflation of apparently very highly significant effects for small differences in PI values (e.g. -0.1 vs 0.1), leading to problems with replicability. Here it is shown on the basis of a well-established statistical model for binary decisions that application of the t-test to PI data implicitly assumes that each fly in a batch is an independent biological replicate in the extremely strict sense that the decision of each individual was interrogated independently of the other flies. Therefore, t-test analysis of PI data obtained with the intrinsically pseudoreplicative group assay is the cause of extremely optimistic P-values, as suggested recently by Bassetto et al. (2023) on the basis of effect size considerations for proportions. Thus, rather than using inferential statistics, PI data should be assessed on the basis of effect size. A more fundamental problem with the batch PI value is the uncertainty in whether it measures the mean individual preference and if so, with what precision, given the simple readout? This aspect is illustrated here by modelling distributions of flies in the T-maze, which suggest that the effective precision of the PI value is clearly worse than the nominal precision of +/- 0.01 for 100 flies, so that the batch PI can only serve as a rough indicator of group tendencies.

## Introduction

The binary choice chamber (T-maze) assay is a standard behavioural screen in Drosophila research to test mutants in terms of sensory discrimination skills or learning aspects, such as memory consolidation and decay (Tempel et al. 1983). Typically, a batch (cohort, group) of ca. 100 flies of a given genotype is tested all at the same time, and as with most screening methods, the readout is not sophisticated, but based on counting the number #*A* of flies in the test arm exposed to the stimulus versus the number #*B* of flies in the control tube, both at the end of the experiment, e.g., 2 minutes after the T-port was opened for the flies to enter the T-maze. The difference in fly count between tube A and B, normalized by the total, defines the batch preference index (PI)

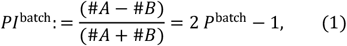

where *P*^batch^ = #*A*/(#*A* + #*B*) is the proportion of the flies found in tube A. For statistical data analysis of sample *PI*^batch^ values, the Student t-test was recommended in protocols papers (Krashes & Waddell 2011a,b), giving reference to the invaluable compendium of biostatistics by Zar. Looking into the book (Zar 1996, p.553, Section 23.11), we find the proposition “If the two proportions to be compared are the means from two sets of proportions, then the two-sample t-test could be used, in which case the data should be transformed … preferably using the arcsine transformation”. The transformation is necessary to account for the unusual binomial error distribution for a proportion, having maximum variance at 50% and zero variance at the strict boundaries of 0% to 100%. As obvious from Eq. (1), the PI is only a linear function of the proportion and thus still has the binomial error structure. With a few exceptions (e.g., MacWilliams et al. 2018), most researchers apply the t-test directly to the PI values of the sample without applying a non-linear transformation. The bigger problem is that the advocates of the t-test apparently were not aware of Zar (1996) making his suggestion on the understanding that the two proportions to be compared with the t-test were obtained from many independent observations of a binary random variate, in other words, each individual animal would need to be tested individually and chose independently between A and B. For example, Zar’s procedure would apply to an opinion poll comprising a sample of 1000 randomly selected participants from ten regions (ten sets) with 100 randomly selected persons each, always with each individual asked to give a yes-no answer secretly in a polling booth. Then to statistically infer the opinion of the whole population, one may compute the test statistics *t* for the set of 10 subsample proportions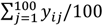 (preferably submitted to the arcsine transformation), with *y*_*ij*_ being 1 for ‘yes’ and 0 for ‘no’ of the *j*-th individual in the *i*-th set, or either simply add up all individual answers across all sets 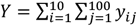, to obtain the mean proportion *Y*/1000 to be analyzed with the exact binomial test, available in a decent software for statistical analysis. Clearly, from a modern perspective, application of the t-test to such cases is obsolete, as the proposition of Zar dates back to times when most statisticians performed manual calculations of test statistics and looked up the extreme values for the test statistics in tables compiled in books. Back then, it may indeed have been simpler to work out the t-test statistics, compared to an exact binomial test. Regardless, if there are *Y*=550 ‘yes’ opinions in total, then the probability of observing a sample with *Y/1000*=55% given no preference in the population (“fifty-fifty”) would be ca. 0.0017 according to the exact binomial test. In a nutshell, the P-value is 0.0017, and we can reject the null hypothesis of no preference in the population at a significance level of *α*=0.002. A different way of putting it is to say that the 1 − *α* = 99.8% confidence interval [50.07%;59.9%] for the preference in the whole population does just not include 50%.

But what if the poll workers take a shortcut and now shout to 10 crowds of 100 people each, asking them to raise their hands if they support a proposition or not? How much confidence would we now have in a mean proportion of 55% and how meaningful would it be? In particular, would we be inclined to believe a statistical test that yields a P-value of 0.0017 for the shortcut scenario? And what if the poll statistician did not know of the shortcut and assumed 1000 independent observations in a secret poll?

With this in mind, let us now consider the action of the t-test when applied to sample *PI*^batch^ values for the case of no pronounced preferences for a stimulus (i.e. *P*^batch^ values close to 50%). Based on the *t*-test, Gegear et al. (2008) obtained 4-star significant P-values (below 0.0001 significance level) for small proportion contrasts, such as 45.6% (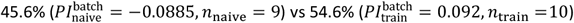 between naïve and trained flies, tested in batches comprising 100 to 150 flies each (see Fig. 1b of Gegear et al. 2008, white eyed *w*;Canton-S). The proportions 45.6% vs 54.6% translate into an odds ratio of OR=1.44, which is a small effect size according to conventional classifications of effect sizes (e.g., Chen et al. 2010). Thus, the 4-star significant P-values are in stark contrast to the small effect size. The sensory task in Gegear’s assay given to starved flies was to discriminate between the background magnetic field of the Earth and a ca. 10 times stronger magnetic field (5 Gauss or 500 μT), where training consisted in the one-time exposure of starved naïve flies to the stronger field when given the opportunity to feed on sucrose for ca. 2 minutes. No furthering experiments with a different technique were provided to back up the conclusions drawn exclusively from T-maze data. We can thus ask ourselves again how much confidence do we have in such small proportion contrasts representing biologically meaningful effects? And of course, does such a result stand up to the gold standard of independent replication?

**Figure 1:**
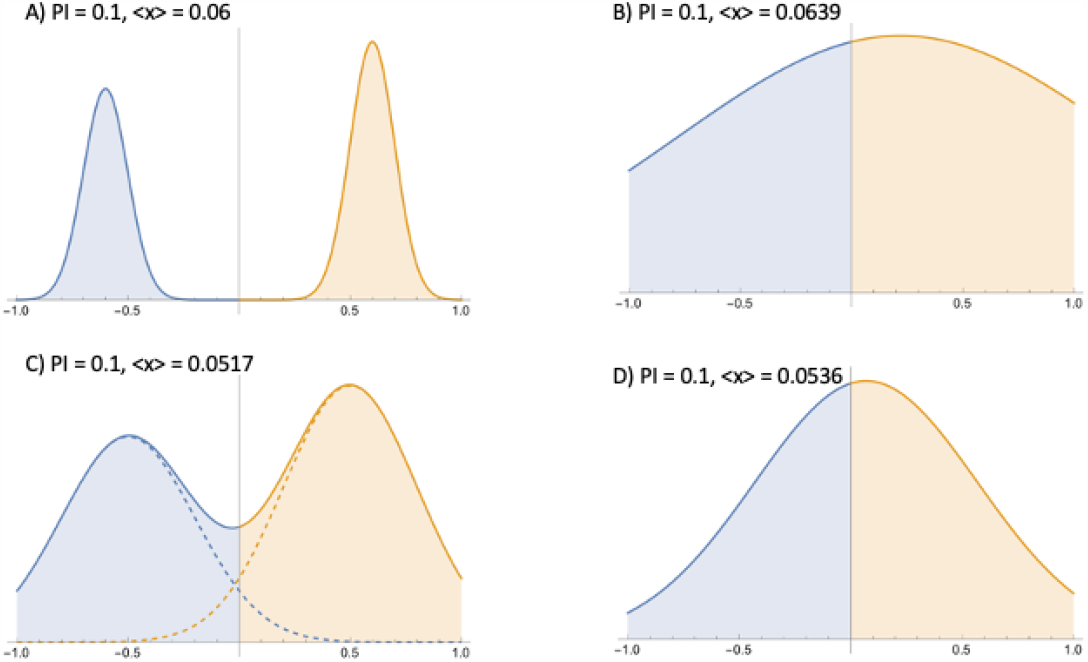
Four different distributions of flies in the T-maze, each having a PI of 0.1 (weak preference toward the right), but all indicate a different weighted mean preference <x>. A) ideal (separable bimodal), C) overlapping bimodal, B) and D) monomodal with larger dispersion in B) compared to D). The ideal bimodal distribution A with two clearly separated partial distributions is the only distribution from which a meaningful quantitative proxy of the group preference can be derived with a defined precision. The <x> values of B, C, and D differ by +7%, -14%, and -11% from that of the ideal distribution. Note that distribution A) has the lowest readout noise, but even so, the problem of pseudoreplication remains.

The work of Gegear et al. (2008) was criticized early on by Mora et al. (2009) who objected amongst other things that

i. the (conditioned) behavior may not be independent of any probabilities of aggregation,
ii. the effect(s) of social facilitation in large groups is/are unknown, and
iii. the appropriate choice of sampling unit for replication (i.e., whole group versus individual animals) may not be clear.

In other words, Mora et al. (2009) identified the group assay as pseudoreplicative (point i), which makes it very hard to find an appropriate statistical framework for undoing pseudoreplication mathematically (point iii). There are different types of pseudoreplication (see excellent review articles by Hurlburt 1984 and Forstmeier et al. 2017), but they all have in common that they produce clustered data, overestimate the actual statistical degrees of freedom, and consequently yield overly optimistic P-values. Forstmeier et al. (2017) considers incorrect P-values due to unrecognized pseudoreplication as a major contributor to the “reproducibility crisis” in science.

Indeed, in a well-controlled, large-scale replication attempt, Bassetto et al. (2023) failed to see any magnetic-field effects on fly behaviour, neither in naïve nor in trained flies, despite having used the same T-maze design and magnetic field conditions as in the original report of Gegear et al. (2008). Further, on the basis of effect-size considerations and power calculations for proportions, Bassetto et al. (2023) found that 4-star significance for small proportion contrasts can only be obtained when each fly in each batch would be counted as an independent biological replicate, so that *n* = 10 batches with *n* = 100 flies each, would count falsely as *n* × *m* = 1000 independent biological replicates. They thus correctly identified the t-test as the culprit in the original work, responsible for the inflation of apparent highly significant effects. Proponents of the t-test might object that their statistical analysis is based on *n* independent sets of flies (sample size: *n*; statistical degrees of freedom: *n* − 1). Indeed, it may not be straightforward to see where the t-test implicitly includes *m* when passing just *n* PI values to their t-test interface. Obviously, *m* enters in the form of #A and #B counts (Eq. 1), and it is exactly here that the implicit assumption is made that each individual in the crowd counts as an independent unit of replication.

In the Appendix included at the end of his paper, a statistical proof is provided that the one-sample Student’s *t*-test applied to *n* batches, with *m* flies each, implicitly assumes *n* × *m* independent observations. It thus implicitly assumes the secret-poll scenario when in fact the data collection was the pseudoreplicative crowd scenario. The resulting test statistics *t*_*n*_ for the secret-poll scenario in the limit of large *m* is obtained as

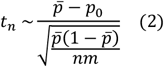

(Eq. A5 in Appendix), where 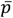 is the mean proportion and *p*_0_ is the proportion under the null hypothesis (*p*_0_=1/2 for zero preference). Eq. (2) is approximately equal to the test statistics *Z* determined from many independent observations (large sample size *nm*),

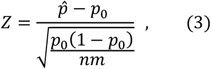

(Eq. 23.42 in Zar 1996), where 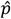 is the estimate for the population proportion, which is equal to the mean proportion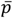 obtained by averaging over the *n* subsamples. When the subsample size is not the same for all subsamples, then the equality 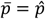 is achieved by taking the weighted mean of subsample proportions, each weighted by the subsample size. The difference between 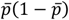 from Eq. (2) and *p*_0_(1 − *p*_0_) is less than 10% for 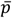 between 0.35 and 0.65, but most importantly, both test statistics are proportional to 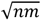, therefore we can say that for large *m, t*_*n*_ approximates *Z*, and note that generally *t*_*n*_ converges to *Z* for large *n* (as the Student t-distribution converges against the normal distribution for large *n)*. The result above (Eq. 2) also confirms the proposition of Zar (1996), who certainly did not have a pseudoreplicative ‘crowd’ scenarios like the Drosophila T-maze tests in mind when proposing this approach to analyze proportions from a large number of independent observations.

To summarize, the t-test applied to data derived from batches of flies implicitly assumes the secret poll scenario, considering each single fly in a batch of *m* flies as independent replication unit and thereby inflates the statistical degrees of freedom by a factor of *m*. This in turns boosts the test statistics by a factor of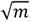, thereby making the test statistics look extreme, leading to highly exaggerated P-values. Put simply, the t-test and (similarly ANOVA for more than two conditions) is a fundamentally wrong test for the problem at hand. Without providing a formal proof here, non-parametric rank tests such as the Wilcoxon Rank Sum test, a.k.a Mann-Whitney U-Test are also inappropriate, as can be already seen from their action on such data, again yielding exaggerated P-values for small proportion contrasts. Trying out various other tests in the pursuit of significance, a sad practice referred to as p-value hacking, is no option either. The problem is pseudoreplication, and this has to be recognized at the outset if it was not openly declared by the experimenter.

This of course begs the question how to statistically correctly analyze proportion data based on group assays such as the Drosophila T-maze assay?

If a statistical test is needed, the test statistics must be corrected for pseudoreplicates. A strict approach has been suggested by Bassetto et al. (2023), see Eq. (A9-A11) here, which divides the original pseudoreplicative test statistics by 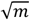, thereby undoing the artificial inflation. It is clear that the ensuing test would then yield a conservative P-value, so that a meaningful conclusion could be drawn already from a P-value below a significance level of 0.1. This level would be considered as marginally significant for a well-defined problem for which a powerful test exists, but pseudoreplicative data from T-maze experiments present an ill-posed statistical problem and it is not clear if a powerful test exists at all. Alternatively, one can follow Hurlburt’s (1984) advice to statisticians “to not let experimenters sweet-talk you into condoning a statistical analysis where accuracy would be better served by not applying inferential statistics at all”. This implies effect-size considerations, which for T-maze data should be based on Cohen’s effect size *h* for proportions, as urged in Bassetto et al. (2023). Such considerations could forestall wrong statistical inference and irreproducible effects.

### Precision considerations

Irrespective of the pseudoreplication problem, *PI*^batch^ is generally treated as if it were a measurement variable with a very high precision. However, a fundamental problem with the *PI*^batch^ is that it purports to “measure” the mean decision of *m* flies with a numerical precision of ±0.01 (for a batch of *m* =100 flies). The problem is that *PI*^batch^ is not sensitive to the degree of individual decidedness. Some individuals may make a firm decision, moving quickly into a tube and staying there, others may turn back and continue their search for food in the opposite tube, while some others may be random walkers and just happen to be trapped anywhere when the preset duration of the experiment is over. All three types enter with the same weight in the binary counting scheme. When looking at video recordings of T-maze data (e.g., https://www.youtube.com/watch?v=-6kt2T4T0Ys ), one can clearly see sustained fly traffic in the T-piece between the two tubes, which prompts the question of whether or not there is a stable end point at which no more left-right fluctuations occur.

To address the problem, let *x* the coordinate of the T-maze extending from left to right (−1 ≤ *x* ≤ 1) with *x* = 0 defining the neutral position (inlet) in the middle. Let *x*_*j*_ the position of fly *j* in the T-maze, then the center of mass of the batch is given by

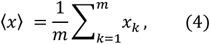

which can be regarded as the weighted mean decision. In contrast,

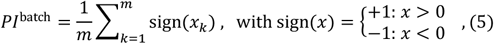

simply binarizes the positions, regardless of the actual distance travelled. Therefore, there will be many different realizations of ⟨*x*⟩ values that have the same *PI*^batch^ value, so there is no one-to-one mapping between *PI*^batch^ and the meaningful ⟨*x*⟩ parameter. Without *PI*^batch^ and ⟨*x*⟩ being connected by a monotonic function (over the value range of interest), one cannot hope to use *PI*^batch^ values as a faithful proxy of the group decision ⟨*x*⟩, notwithstanding the purported precision figure of 0.01.

This is illustrated in Fig. 1, with an ideal, separable distribution of flies (Fig 1a), as opposed to more realistic distributions with overlap (“traffic”, see Fig. 1b,c,d). All four distributions have the same integral preference index (0.1, slight preference toward right), here defined as the normalized difference of the cumulative distribution cdf(*x*) between right and left side,

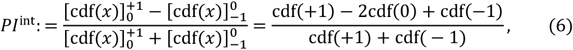

where 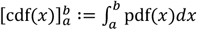 pdf(*x*)*dx*, where pdf(*x*) is the probability distribution function, yet the weighted mean preference,

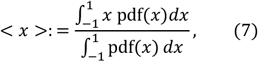

Is different in each case. This simple example shows how much uncertainty the PI has as a proxy parameter if a stable end point does not exist, with a stable end point defined as two clearly separated monomodal distributions as in Fig. 1a. In particular, it is only for the stable end point (Fig. 1a) that the PI has a defined numerical precision of ±1/*m*, while traffic (i.e. overlap in the other distributions Fig. 1b-d) debilitates this precision figure. Thus, to estimate the readout noise, a photo is needed to document the distribution of flies in the setup when the trial time is up, just before the T-port is locked and flies are trapped in either testing tube. From the photo, ⟨*x*⟩ can be determined according to Eq. (4) and be compared with the PI obtained from the binary counting scheme (Eq. 5). Repeating this procedure for several batches (sample size *n*) then allows one to determine the degree of correlation between PI and ⟨*x*⟩ by regression analysis (e.g. linear model( PI ∼ ⟨*x*⟩ ), with PI as response variable and ⟨*x*⟩ as predictor variable. For each observation (PI, ⟨*x*⟩)_*i*_ with (*i* = 1, …, *n*), this procedure yields a predicted *PI* value, to be subtracted from the observed *PI* value. The difference could serve as a precision figure for each PI. This would help quantify the readout noise, but does of course not mitigate the intrinsic problem of pseudoreplication.

## Conclusions

The pseudoreplicative nature of the T-maze assay leads to an inflation of statistical effects when using the t-test. This was already concluded by Bassetto et al. (2023) on the basis of effect size and power calculations and emerges here naturally from a statistical model. Therefore, as pointed out by Bassetto et al. (2023), T-maze data should always be judged in terms of Cohen’s effect size *h* for proportions, with *h* being the difference between two arcsine-transformed mean proportion estimates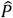,

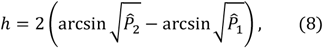

(max(*h*) = 2), or in terms of an odds ratio,

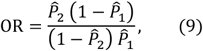

with 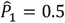 for the one-sample case against no preference. According to the guidelines of the American Psychological Association, confidence intervals for the effect sizes should be reported to see if *h* = 0 or OR = 1 is included, and this can be computed conveniently with resampling techniques (bootstrap), see Banjanovic & Osborne (2019) for details.

The point of using several sets of flies in a given sample (condition) is to obtain a robust estimate of 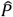 for effect size calculations. Importantly, the within-sample dispersion of preference indices is irrelevant for proportion statistics and effect sizes for proportions, but exactly that dispersion measure is used for t-test and ANOVA-based inferential statistics and effect size calculations. Therefore, the within-sample dispersion (e.g., in form of standard error of the mean, s.e.m.) should not even be reported for proportions and for preference indexes. Also, for t-test and ANOVA, it is assumed that the within-sample dispersion describes the sampling noise, due to taking random samples from a population. There is no room for readout noise, which is considered negligible compared to the sampling noise. However, the readout noise of the T-maze assay remains to be assessed. As suggested above, this requires photo documentation of the distribution of flies in the T-maze at the end of the experiment.

The intrinsic uncertainty of whether *PI*^batch^ values in the range -0.1 to 0.1 (and possibly beyond) even measures the group averaged decision remains a fundamental limitation. Thus, when wishing to adhere to the batch assay, the readout needs to be improved in order to increase the sensitivity of the assay. An obvious possibility would be to video the flies and track their positions, as done in the negative geotaxis assays (gravity, FlyVac) presented in Bassetto et al. (2023). This would allow one to determine the weighted mean decision over time and validate the stability of the end point, from which the precision of the PI could be obtained. Also, video analysis would allow one to estimate diffusive properties of a batch, which could be fitted to hidden Markov processes or models of random walks, which in turn would provide insights in group facilitation or distraction.

### Practical advice

It is clear that one cannot hope to obtain accurate P-values from pseudoreplicative experiments (Hurlburt 1984). However, the conservative approach proposed in Bassetto et al. (2023) provides a means to make sure that statistically significant effects seen in T-maze data are almost certainly real and the inferred effects highly likely to be reproducible. The test for equality of two sample proportions is based on a general linear model with binomial error structure, with an interface of the form

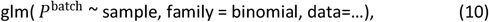

with the factor variable sample (e.g., factor levels naïve and trained). But note, that the following formula,

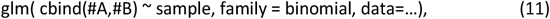

where #A,#B as in Eq. (1) would assume each fly as an independent degree of freedom (pseudoreplication). Although Eq. (11) casts a warning for non-integer values, it is formally the correct way of avoiding pseudoreplication. Put differently, when the command in Eq. (10) is executed, the machine has no way of knowing that #A,#B are counts from a pseudoreplicative assay.

## Acknowledgements

The author is grateful to Marco Bassetto, Dmitry Kobylkov, Henrik Mouritsen, and Peter Hore for fruitful and lively discussions about fly behaviour, statistics, magnetic conditioning, and scientific standards. The author would like to thank Henrik Mouritsen and Peter Hore for valuable comments on this manuscript.

## Appendix

Here we will prove first that the t-test statistics *t* for *n* sets of *m* independent observation is equivalent to the test statistics *Z* for the proportion for *n* × *m* independent observation. The implication is that by applying a Student *t*-test to the values of *n* batches of *m* flies each, one implicitly assumes an ideal experiment with *n* times *m* independent flies, and therefore obtains pseudoreplicative p-values. A correction of the test statistics for pseudoreplication is derived in the second part.

### Definitions

Let us define an ideal experiment where *m individual flies are tested one at a time*. For these *m* independent trials, let *m*^+^ be the number of successful outcomes (stimulus arm) and *m*^−^ the number of unsuccessful outcomes (control arm). The corresponding 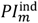 then is defined as

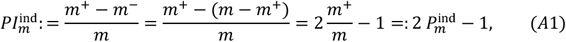

where 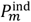 is the proportion of successful outcomes. Eq. (A1) links the two definitions via 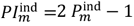, which superficially resembles *PI*^batch^: = 2 *P*^batch^ − 1 from Eq. (1) in the main text. However, this resemblance is deceptive, because *PI*^batch^ is based on group behaviour while 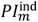 is based on independent trials, whence the different superscripts here.

#### Assumptions

Let us now assume a Bernoulli distribution such that each fly at the entry to the T piece has the same probability *p* of selecting one tube (e.g., stimulus port) and a probability of 1 − *p* of selecting the other port (control port), and again that each fly is independent, i.e. tested singly. Then the theorem by de Moivre and Laplace states that the asymptotic distribution (i.e., large *m*) of the quantity

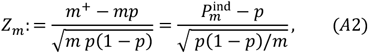

is the standardized normal distribution 𝒩_0,1_(*Z*_*m*_); in particular, for large *m*, 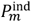 is a normally distributed random variate, with expectation value 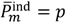 and standard deviation 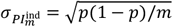 .

Similarly, 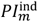 is a normally distributed random variate, too, with expectation value 2*p* − 1 and standard deviation 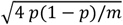, because Var(*c X*) = *c*^2^Var(*X*), where *c* is a constant (here *c* =2) and *X* denotes any random variate. 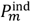 can be used as a diagnostic to test the model, i.e., if the underlying statistical process is indistinguishable from a Bernoulli process. Now for the sake of argument, consider the case where *n* independent sets of experiments are conducted (e.g. by *n* different workers or by 2 different workers on *n/2* different days), each time with a new series of *m* independent flies tested one at a time. Then the test statistics *Z*_*nm*_ for these *N* = *n m* independent trials in the binomial test (single proportion test) for the large sample approximation according to Eq. (A2) is given by

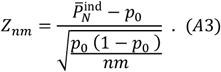

#### Proof of equivalence

If the one-sample t-test is to be used to find out if 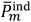, the mean of 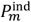 from *n* repeats, differs significantly from the value according to the null hypothesis (here *p*_0_ = 1/2 ), then the t-statistic needs to be considered, given by

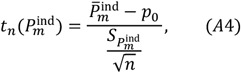

where 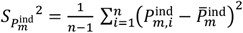 is the sample variance. It is clear that a sum of squared random variates is distributed as 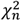 (chi-squared with n degrees of freedom), but making use of the central limit theorem for large *m* (say *m* > 30), the distribution of, 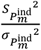 where 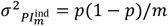 will approximate a normal distribution centered at 1 with scale parameter 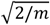, so that we can substitute 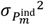 for 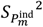 in Eq. (A4) to obtain a statistical approximation of 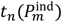 as

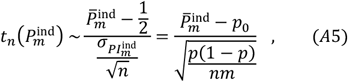

which for large *m* is statistically equivalent to *Z*_*nm*_ (compare r.h.s. of Eq. A5 with Eq. A3). In other words, given an underlying Bernoulli process, the *t*-test statistics for *n* independent sets of *m* ideal experiments is equivalent to the test statistics *Z* for the proportion of flies that in *n* × *m* independent ideal experiments chose the desired port.

#### Pseudoreplication

Now we turn to batches of flies, where the assumption of independence is violated. Applying the 1-sample *t*-test to *n* batches of flies against the null hypothesis of no effect at the population level ( *ℋ*_0_: μ = 0), consists in computing the following *t*-statistic

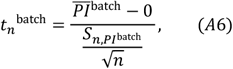

which because of *PI*^batch^: = 2 *P*^batch^ − 1 (Eq. 1) is the same as

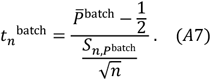

with *ℋ*_0_: *p*_0_ = 1/2. Eq. (A7) can now be compared with the t-test statistics for the ideal experiment with *n* times *m* flies tested one at a time derived from Eq. (A4), i.e,

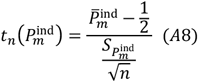

It is clear that Eq. (A8) can be equated to Eq. (A7) if and only if *t*_*n*_^batch^ is a reliable proxy of 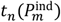 and that each fly in a batch is an independent biological replicate, with is most questionable in a pseudoreplicative assay.

From power calculations, Bassetto et al. 2023 used the following expression for the test statistics Z for *n* batches under the assumption the independent unit of replication was a batch of flies as

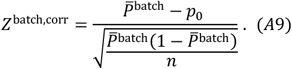

Comparing Eq. (A9) with Eq. (A3), where *Z*_*nm*_ was based on *N* = *nm* independent biological replicates, we have

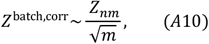

provided that 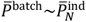, i.e. provided that 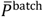 approximates 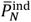 .

By analogy with Eq. (A10), the correction of 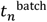 for pseudoreplication can be obtained as

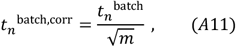

which follows from the statistical equivalence of *Z*_*nm*_ and 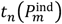 (again, compare r.h.s. of Eq. (A5) with Eq. (A3)). It goes without saying that Eqs. (A10) and (A11) also apply to two-sample tests, provided that batch sizes are roughly similar for the two samples.

In addition to being pseudoreplicative, the test statistics 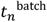 may have little precision because of the unknown precision of *P*^batch^ and thus also of 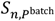, the within-sample variance of *P*^batch^. The latter may underestimate the true dispersion in which case it produces larger (i.e. “more significant”) 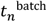 values. Therefore, the rationale behind Eq. (A10) and the approach of Bassetto et al. 2023 is to take the average proportion over *n* batches as an estimate of the population proportion 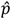, and use the standard error of the proportion, 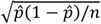, instead of the standard error of the mean, to estimate the confidence intervals.

#### P-values corrected for pseudoreplication

In the proportion test, the “P-value” is the probability of the test statistics *Z*^batch,corr^ under the assumption that the null hypothesis 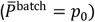 is true,

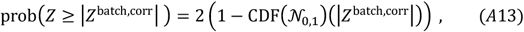

where CDF(𝒩_0,1_)(*x*) denotes the cumulative distribution function of the standardized normal distribution and the factor 2 accounts for the two-sided alternative hypothesis, 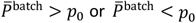 . Similarly, for the t-test, the P-value of the test statistics 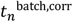 given that the null hypothesis 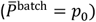 is true, writes as

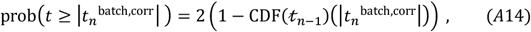

where 𝓉_*n*−1_ is the Student t distribution with *n* − 1 degrees of freedom for a sample size of *n*.

Example:

For a proportion contrast of 45.6% to 54.6% between naïve (*n*_naive_ = 9) and trained flies (*n*_train_ = 10) tested in batches (white eyed *w*;Canton-S in Fig. 1b of Gegear et al. 2008), a P-value of <0.0001 based on the Student t-test was reported in Gegear et al. 2008. From the critical value of *t* for *v* degrees of freedom at significance level *a* under the assumption that the null hypothesis is true,

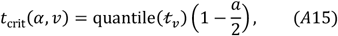

a P-value of (<0.0001) implies an uncorrected two-sample statistics of 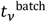 *> t*_crit_(0.0001, 17) = 5.05, where *v* = *n*_naïve_ + *n*_train_ − 2 = 17. In fact, from the published data (*PI*_naïve_ ± s. e. m. = −0.0885 ± 0.023 versus *PI*_train_ ± s. e. m. = 0.092 ± 0.019), *t*_*v*_^batch^ is ca. 6.0. With 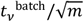 corrected for pseudoreplication (batch size *m*), the corrected P-value comes out as non-significant (ca. 0.5), which is reasonable given the small sample sizes and small effect size (OR=1.44).

It is clear that the 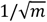 correction of the test statistics (Eq. A10, A11) will diminish the false positive rate at the expense of increasing the false negative rate. As this is a conservative approach, a P-value of 0.1 would now indicate a significant result. In contrast, with the widely used, but uncorrected form of the statistics (Eq. A7), the false positive rate gets out of control, as indicated by 4-star significances for the example in the introduction. Therefore, extreme caution is needed when drawing scientific conclusions only on statistical inference based on the uncorrected Eq. (A7) without further experiments using a different method.

